# PACAP-mediated gating of anxiety-controlling circuits

**DOI:** 10.1101/2023.05.01.539007

**Authors:** Yan Li, Raül Andero, Natalia V. Luchkina, Junghyup Suh, Rachel A. Ross, Bradford B. Lowell, William A. Carlezon, Kerry J. Ressler, Vadim Y. Bolshakov

**Author notes:** Correspondence to: V.Y.B.

## Abstract

Combining the use of *ex vivo* and *in vivo* optogenetics, viral tracing, electrophysiology and behavioral testing, we show that the neuropeptide pituitary adenylate cyclase-activating polypeptide (PACAP) gates anxiety-controlling circuits by differentially affecting synaptic efficacy at projections from the basolateral amygdala (BLA) to two different subdivisions of the dorsal subdivision of the bed nucleus of the stria terminalis (BNST), modifying the signal flow in BLA-ovBNST-adBNST circuits in such a way that adBNST is inhibited. Inhibition of adBNST is translated into the reduced firing probability of adBNST neurons during afferent activation, explaining the anxiety-triggering actions of PACAP in BNST, as inhibition of adBNST is anxiogenic. Our results reveal how innate, fear-related behavioral mechanisms may be controlled by neuropeptides, PACAP specifically, at the level of underlying neural circuits by inducing long-lasting plastic changes in functional interactions between their different structural components.

## Introduction

Mechanisms of behavioral control are determined by the function of underlying neural circuits (Shumyatsky et al., 2005; Janak and Tye, 2015; Basu et al., 2016; Tovote et al., 2016). To understand the logic of circuit-level computations guiding behavior, it is necessary to define how the interactions between components of the circuitry are modulated in order to achieve certain behavioral outcomes. One of the most efficient strategies in the brain for directing information flow is the release of neuropeptides, which have strikingly specific effects on the function of behavior-driving microcircuits Huber et al., 2005; Tye et al. 2011; Viviani et al., 2011; Knobloch et al., 2012; Van den Pol, 2012; Al-Hasani et al., 2015; Tovote et al., 2015). However, synaptic and network-level mechanisms of neuropeptidergic modulation of behavior are far from being completely understood. Here we focus on the mechanisms underlying innate behavioral responses, such as innate fear, which are amenable to analyses at multiple functional levels and might be universal across species, including humans.

Although the circuitry of innate fear-related behaviors is quite extensive, there are certain regions of the brain which have been consistently implicated in the regulation of innate fear, including the amygdala and the bed nucleus of the stria terminalis (BNST) (Pleil et al., 2015; Henckens et al., 2016), both of which are enriched with neuropeptides. The BNST, with its numerous subregions, shows unique diversity in terms of neuronal populations and projections (Bota et al., 2012; Daniel and Rainnie, 2016), and recent studies in humans (Somerville et al., 2010) and animals (Duvarci et al., 2009) point to a major role of the BNST in anxiety-related behavioral manifestations. Specifically, the amygdala may be responsible for an immediate, phasic fear response, whereas the temporally prolonged, sustained fear, or anxiety, may be represented by activity of the BNST (Davis, 1998; Walker et al., 2009; Davis et al., 2010). Whereas many neuropeptides affect neuronal functions (e.g., Viviani et al., 2011), only a handful of them have been linked to control of fear-related behaviors. Pituitary adenylate cyclase-activating polypeptide (PACAP) is particularly interesting in relation to fear mechanisms, as there is evidence that PACAP-mediated signaling regulates fear states (Vaudry et al., 2009; Ressler et al., 2011; Hammack and May, 2015).

Here we report that neuropeptide PACAP gates innate fear circuits by differentially affecting synaptic efficacy at projections from the basolateral amygdala (BLA) to two different subdivisions of the dorsal subdivision of the bed nucleus of the stria terminalis (BNST) (Pleil et al., 2015; Henckens et al., 2016), modifying the signal flow in BLA-ovBNST-adBNST circuits in such a way that adBNST is inhibited. Inhibition of adBNST is translated into reduced firing probability of adBNST neurons, explaining the anxiety-triggering actions of PACAP in BNST, as inhibition of adBNST is anxiogenic. Our results reveal how innate, fear-related behavioral mechanisms may be controlled by neuropeptides at the level of underlying neural circuits by inducing synaptic plasticity-mediated changes in functional interactions between their different structural components.

## Methods

### Animals and surgery

Five to six week old male C57BL/6 mice (Charles River Laboratory) were used for stereotaxic surgery and viral injections. All animals were housed on a 12-hour light cycle (light on at 7 AM) with ad libitum access to food and water. All animal procedures were approved by the Institutional Animal Care and Use Committee (IACUC) at McLean Hospital and conducted in accordance with the National Institutes of Health guidelines. Adeno-associated virus (AAV) carrying channelrhodopsin-2(H143R)-eYFP under control of either CaMKIIα or hSyn promoters, packaged by the Vector Core facility at the University of North Carolina, were used to target the BLA-BNST and PBn-ovBNST projections, respectively. Viral titers were 4.7-6.2 x 10^12^ particles/ml and 4.8 x 10^12^ particles/ml for AAV5-CaMKIIα-ChR2(H134R)-eYFP and AAV5-hSyn-ChR2(134R)-eYFP, respectively. Before surgery, mice were anesthetized with a mixture of ketamine and xylazine (160 mg/kg and 12 mg/kg body weight, respectively). All surgical procedures were previously described in detail (Cho et al., 2013). Briefly, bilateral craniotomy was made to target the basolateral nucleus of the amygdala (BLA) with stereotaxic coordinates: 1.6 mm caudal to bregma, ± 3.2 mm lateral to midline, and 4.3 mm ventral to bregma (Franklin and Paxinos, 2008). Virus was injected into BLA bilaterally (0.5 μl per side) with a Hamilton microsyringe and a syringe pump (Stoelting Co.) at a rate of 0.1 μl/minute. Mice were given a subcutaneous injection of ketoprofen (5 mg/kg, Fort Dodge Animal Health) to reduce postoperative pain. After viral injections, mice were single housed for 5-6 weeks before ex vivo electrophysiological experiments.

### In vitro electrophysiological recordings and photostimulation

Coronal brain slices (300 μm in thickness) containing either the amygdala or the BNST, as well as the PBn (in experiments focusing on PBn-ovBNST projections) were sectioned with a vibratome in ice-cold solution containing (in mM): 252 sucrose, 1.0 CaCl2, 5.0 MgCl2, 2.5 KCl,1.25 NaH2PO4, 26 NaHCO3, 10 glucose, equilibrated with 95% O2 and 5% CO2. Slices were then incubated in oxygenated artificial cerebrospinal fluid (ACSF) containing (in mM): 125 NaCl, 2.5 KCl, 2.5 CaCl2, 1.0 MgSO4, 1.25 NaH2PO4, 26 NaHCO3 and 10 glucose at a room temperature before recordings. The dorsal division of BNST was visualized under the microscope (Carl Zeiss) and defined by structural landmarks of the internal capsule (ic), anterior commissure and the stria terminalis (Dong et al., 2001). Whole-cell recording were obtained from neurons in the oval nucleus of BNST (ovBNST) and/or anterodorsal nucleus of BNST (adBNST) with patch electrodes (3-5 MΩ resistance) containing (in mM): 130 K-gluconate, 5.0 KCl, 2.5 NaCl, 1.0 MgCl2, 0.2 EGTA, 10 HEPES, 2 MgATP, and 0.1 NaGTP (adjusted to pH 7.2 with KOH). For experiments involving photostimulation of PBn-ovBNST projection fibers (Fig. 2, L and M), potassium gluconate was replaced with 131 mM Cs-methane-sulfonate, and the free [Ca]in was buffered at ∼100 nM with 5 mM EGTA/1.97 mM CaCl2 in the pipette solution (Riccio et al., 2009). Neurobiotin (6 mM, Vector Labs) was added to the internal pipette solution to locate and reconstruct the recorded neurons in subdivisions of BNST after experiments. Whole-cell recordings were performed at 30-32°C with EPC-9 amplifier and Pulse 8.8 software (Heka Elektronik). Currents were filtered at 1 kHz and digitized at 5 kHz. After recordings, slices were placed in phosphate buffered saline (PBS) containing 4% paraformaldehyde and kept in a refrigerator for the subsequent histological processing. Synaptic responses were induced by photostimulation of ChR2-expressing BLA fibers in BNST through 40X water-immersion objective (IR-Achroplan, Carl Zeiss) with the blue light-emitting diode (LED) (Thorlabs; excitation wavelength 470 nm; 5-ms pulses). Illumination area was 0.28 mm2, and, it was centered at the soma of recorded neurons. Light power was measured at 470 nm with a power meter, (Coherent) which was placed under the objective, and light power density, expressed as mW/mm2, was calculated, dividing light power by illumination area (Cho et al., 2013). Pharmacological reagents used in electrophysiological experiments, including NBQX, D-AP5, bicuculline (all from R&D Systems), tetrodotoxin (from Abcam) and 4-aminopyridine (SIGMA), were prepared as stock solutions in water at 500 to 5000-fold concentrations, and stored at -20°C. Stock solutions of PACAP38 (from BACHEM) and PACAP6-38 (from Abcam) were prepared in standard ASCF at a stock concentration of 0.25 mM and stored at -20°C. Neurobiotin was obtained from Vector Labs and also stored at -20°C.

**Figure 1.**
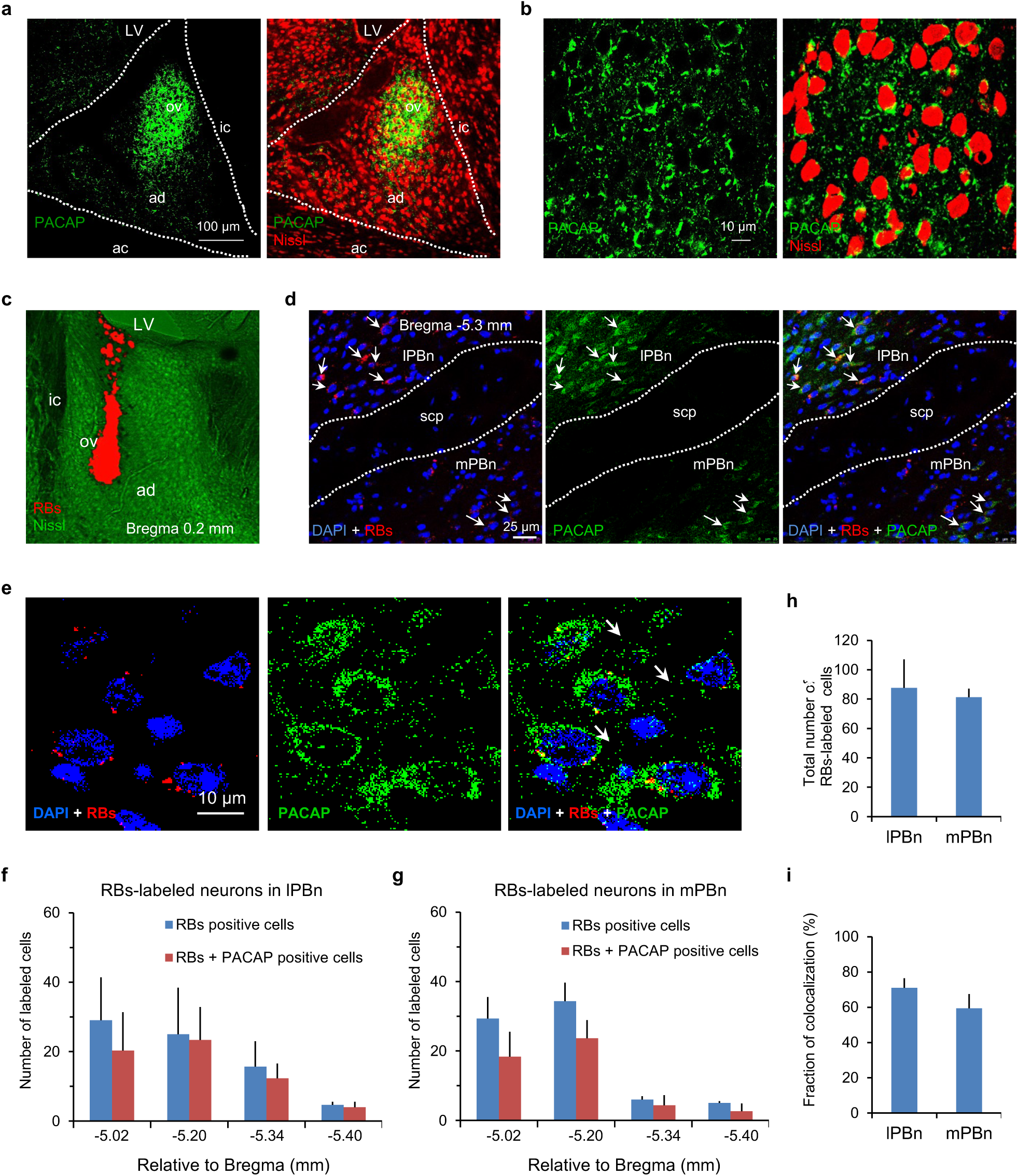
PACAP-containing PBn neurons project to ovBNST. **a**, Left, PACAP immuno-positive fibers were nearly exclusively observed in ovBNST (labeled green); ov, ovBNST; ad, adBNST; ac, anterior commissure; ic, internal capsule; LV, lateral ventricle. Right, fluorescent Nissl staining (labeled red) was combined with immunostaining for PACAP. **b**, High magnification confocal images of PACAP immunostaining in BNST (left), and fluorescent Nissl staining combined with immunostaining for PACAP (right). PACAP immunopositive fibers surrounded somas of Nissl stained neurons in ovBNST. **c**, A microscopic image showing the injection site of retrograde tracer (RBs, red) and fluorescent Nissl staining (green) in the ovBNST. **d**, Left, an image showing RB-labeled cells (red RB puncta can be seen around DAPI-stained blue neuronal nuclei) in PBn (arrowheads). lPBn, lateral parabrachial nucleus; mPBn, medial parabrachial nucleus; scp, superior cerebellar peduncle. Middle, PACAP immunopositive cells (green) in PBn. Right, superimposed images from left and middle. Results of the quantitative analysis showing proportions of double-labeled PBn neurons (RB-containing cells which were also PACAP-immunopositive) are below. **e**, High magnification confocal images showing RB-labeled cells in PBn (left), PACAP immunostaining (middle), and superimposed images of RB and PACAP labeling (right). **f**, Total numbers of RBs-labeled neurons and the RBs-labeled neurons colocalized with PACAP immunoreactivity in the lateral division of parabrachial nucleus (lPBn) in single coronal sections (50 μm in thickness) at different section levels relative to Bregma (averaged data from three independent experiments). **g**, Similar to (**f**) but for the medial division of parabrachial nucleus (mPBn) in same coronal sections. **h**, Total numbers (mean ± SEM) of RBs-labeled neurons in lPBn and mPBn, respectively (*P* = 0.71 for lPBn versus mPBn, Student’s unpaired two-tailed t-test). **i**, Fraction (mean ± SEM) of RBs-labeled neurons which were also immunopositive for PACAP in lPBn and mPBn, respectively (*P* = 0.16 for lPBn versus mPBn, Student’s unpaired two-tailed t-test). See Methods for the technical description of retrograde tracing experiments in combination with immunochemistry for PACAP. Data are mean ± s.e.m.

**Figure 2.**
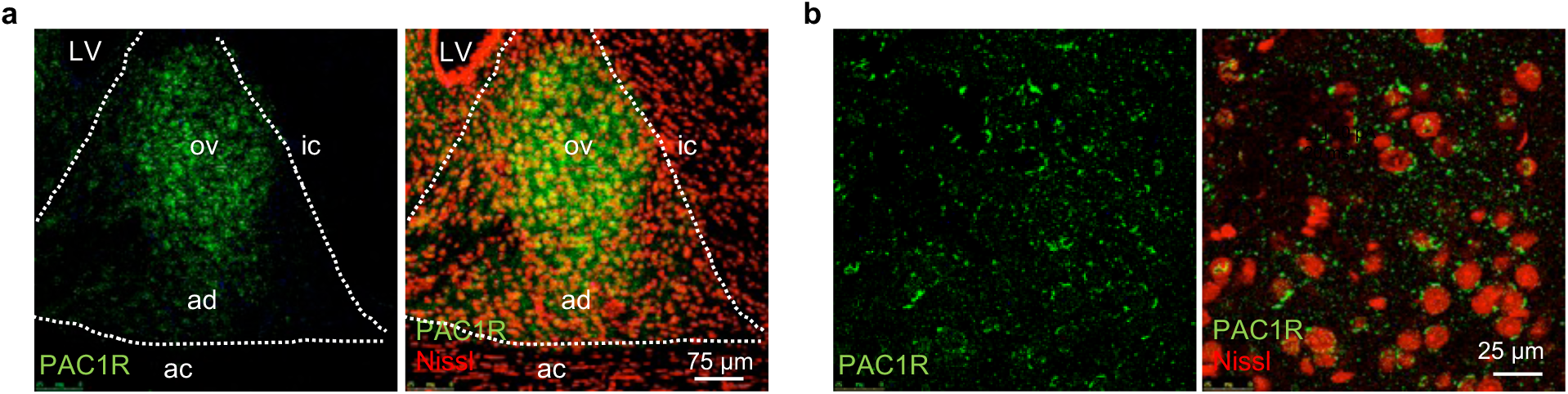
PAC1 receptors are expressed in ovBNST. **a,** Left, PAC1 receptor immunoreactivity was detected in ovBNST (green). Right, Fluorescent Nissl staining (labeled red) was superimposed with immunostaining for PAC1 receptor (green). **b**, Higher magnification images of immunostainings shown in **a**.

### Histology and microscopy

Brain slices were washed in PBS three times, for 5 minutes each time. Slices containing the amygdala or PBn (ChR2 injection sites) and the BNST (projection site) were incubated in 0.1% Triton PBS containing Neurotrace Fluorescent Nissl Stain (Molecular Probes; 615 nm emission wavelength), that was diluted ∼40 fold, for an hour at room temperature, and, then washed again in PBS three times for 20 minutes each time. Slices containing neurobiotin injected cells were incubated in 0.1% Triton PBS containing streptavidin Alexa 568 conjugate (20 μg/ml, Molecular Probes) for two hours at room temperature, and then washed three times in PBS, 20 minutes each time. Finally, slices were mounted on gelatinized slides, dehydrated and coversliped. Vectashield mounting medium (Vector Laboratory) was applied to slices to prevent fluorescence fading. Leica TCS SP8 confocal microscope (Leica) or Zeiss Axioskop 2 fluorescent microscope (Carl Zeiss) were used to capture images. Imaging data were processed and analyzed with ImageJ software (NIH). For each experimental animal, an injection site was verified by visualizing ChR2-eYFP expression. Mistargeted mice were excluded from the analysis.

### Guide cannula implantation for *in vivo* optogenetic behavioral tests

To optogenetically activate projecting fibers originating in parabrachial nucleus (PBn) in ovBNST during behavioral tests, we expressed ChR2 under control of human synapsin promoter (hSyn, pan-neuronal promoter), injecting AAV5-hSyn-ChR2-eYFP (0.5 μl) into the PBn unilaterally. Stereotaxic coordinates for PBn injections were: 5.3 mm caudal to bregma, ± 1.2 mm lateral to midline and 3.5 mm ventral to bregma. On the side of the viral infusion, a second craniotomy was made targeting the ovBNST with stereotaxic coordinates: 0.2 mm rostral to bregma, ±1.0 mm lateral to midline and 3.8 mm ventral to bregma. The guide cannula (33GAC315GS-4/SPC, Plastics One) for photostimulation was implanted into the skull and fixed by dental acrylic and jeweler’s screws (Plastics One). Mice were randomly assigned in respect to the side of viral injections and cannula implantations. Behavioral tests were performed 10 weeks after viral injections to achieve sufficient functional expression of ChR2 in terminal parts of PBn projecting fibers in ovBNST (Kim et al., 2013). After behavioral tests, mice were transcardially perfused with cold PBS containing 4% paraformaldehyde. Brains were removed and postfixed in the same fixating solution overnight in a refrigerator. Brains were immersed in PBS containing 30% sucrose and sectioned at thickness of 50-100 μm on a microtome. The location of the cannula placement was verified under the Zeiss Axioskop 2 plus fluorescent microscope.

### Behavioral testing and in vivo optogenetic experiments

Before the behavioral test, a fiberoptic cord (0.39 NA, 200 μm diameter, Thorlabs) was inserted into the implanted guide cannula. Mice were allowed to recover at home cage from this handling for 1–5 minutes. Diode-pumped solid-state (DPSS) laser, coupled with 20 MHz Waveform Generator (Agilent) was used to emit light (wavelength 473 nm) for photostimulation, delivered through the fiberoptic cord. Parameters of delivered blue laser light pulses were: 5-ms-long pulses, 30 Hz, 4-5 mW in intensity (measured at the tip of the fiberoptic, 127.4-159.2 mW/mm2). Quatricide PV-15 (Pharmacal Research Labs) was used to clean EPM arms between the tests. To provide evidence that photostimulation of ChR2-expressing PBn fibers in ovBNST was efficient, a group of mice received optogenetic stimulation and their brains were fixed shortly after that. We performed immunohistochemistry for c-fos (activity-dependent neuronal marker) expression on this tissue and found that photostimulation triggered c-fos expression in photostimulated area, confirming that blue light stimulation in vivo resulted in activation of neurons in ovBNST. In behavioral experiments, the elevated plus maze was made of plastic and consisted of two gray open arms (30 x 5 cm) and two gray enclosed arms (30 x 5 x 30 cm) extending from a central platform (5 x 5 x 5 cm) at 90 degrees in the form of a plus. Arms of the same type faced each other. The maze was placed 80 cm above the floor. Mice were individually placed in the center, with the head facing to the closed arm. The elevated plus maze test consisted of a 20-min session divided into 5-min epochs: the pre-stimulation light-off epoch, the light-on epoch and the poststimulation light-off epochs, in order (off-on-off epochs). During the 5-minute light-on epoch, a prolonged photostimulation was delivered unilaterally (see above for details of photostimulation). Behaving mice were videotaped and behavior was analyzed by an experimenter blind to the experimental group. We quantified the number of open and closed arm entries and time spent in the open and closed arms. The open field (OF) apparatus was made of white plastic board in size of 45 x 45 x 30 cm (L x W x H). Mouse was placed in the corner of a brightly lit (100 lux) open field. Analogically to the elevated plus+maze test, OF test consisted of a 20-min session divided into 5-min epochs: the pre-stimulation light-off epoch, the light-on epoch followed by two poststimulation light-off epoch, in order (off-on-off-off epochs). In the OF experiments, photostimulation was delivered identically to the EPM tests. We quantified the number of entries and time spent in the center (34 x 34 cm).

### Immunohistochemistry

Mice were deeply anesthetized and perfused transcardially with ice-cold PBS containing 4% paraformaldehyde. After post-fixation in the same fixative at 4°C overnight, the brains were transferred to the PBS containing 30% sucrose until the brains sank. Coronal sections (50 μm) were cut with a vibratome and collected in PBS. The sections were rinsed in PBS three times before and after incubation in PBS containing 0.3% H2O2 for 30 minutes to quench endogenous peroxidase. The sections were then incubated in serum blocking solution (5% donkey serum albumin in 0.1% Triton PBS) containing the primary polyclonal rabbit anti-PACAP antibody (1: 500 dilution, Peninsula Laboratories) or anti-PAC1 receptor (1:50 dilution, LifeSpan BioSciences, Inc.) at 4°C overnight. Subsequently, the sections were washed with PBS and incubated in PBS containing Alexa Fluor® 488 donkey anti-rabbit secondary antibody (1:200, Invitrogen/Molecular Probes) for 2 hr at room temperature. Finally, the sections were rinsed with PBS, and mounted on gelatin-coated slides with mounting medium containing 4’,6-diamidino-2-phenylindole (DAPI).

### Retrograde tracing

To identify the source of PACAPergic fibers in ovBNST, we used fluorescent latex microspheres (RetroBeads, RBs, excitation wavelength 530 nm and emission wavelength 590 nm, Lumafluor) as a retrograde tracer for localized injections into the ovBNST. Retrograde tracer solution containing RBs was diluted to 1:4 in ACSF and stored at 4°C until use. Mice were anesthetized with ketamine and xylazine prior to stereotaxic surgery. RBs were bilaterally injected into the ovBNST with a micro-syringe pump from borosilicate glass capillaries with a tip size of 30-40 μm. The volume of RBs injected to the ovBNST was 0.2 μl, and they were injected at the rate of 0.1 μl per minute. Coordinates for targeting the ovBNST were: 0.2 mm rostral to bregma, ±1.2 mm lateral to midline, and 3.7 mm ventral to the bregma. Mice were given subcutaneous injections of 0.1 ml ketoprofen to reduce pain after surgery. Three weeks after injection of retrograde tracer, we prepared coronal brain sections with a vibratome and fix them with 4 % paraformaldehyde. We then quantified the number of retrogradely-labeled (RBcontaining) neurons colocalized with PACAP immunoreactivity in the parabrachial nucleus. One in every three slides (50 μm in thickness) of consecutive sections of the brain stem was selected for immunohistochemical staining. A confocal microscope, Leica TCS SP8, was used to sequentially acquire signals of DAPI, PACAP immunoreactivity and RBs under three different laser settings (405, 488 and 532 nm, respectively). Brain sections containing PBn were scanned under a 40x/1.3 oil immersion objective lens in 1 μm thin optical planes (290.6 μm × 290.6 μm) for the depth of 20-40 μm. The RBs-labeled cells were usually distributed over 4-5 sections along the rostrocaudal axis of PBn. Images captured with different fluorescent channels were merged using ImageJ software (NIH). RBs-labeled neurons were defined as cells with multiple red puncta in their cytoplasm (red channel) colocalized with blue DAPI, staining nucleus (blue channel). RBs-labeled neurons were further examined for PACAP immunoreactivity (green) in green channel. Total numbers of RBs-labeled neurons and neurons which were both RBs-labeled and PACAP immunopositive were calculated in each section for both subdivisions of PBn. The number of neurons with colocalized RBs labeling and PACAP immunoreactivity of the total number of RBs labeled neurons provided a fraction of PBn-ovBNST projection neurons which were PACAP positive (Fig. 1). Mice mistargeted during retrograde tracer injections were excluded from the analysis.

### Statistical analysis

Data are presented as mean ± SEM. For statistical comparisons, we used Student’s t-test or ANOVA with Bonferroni’s simultaneous multiple comparisons. Statistical analysis was performed with Minitab16 software (Minitab). P < 0.05 was considered statistically significant.

## Results

### PACAP-containing parabrachial nucleus neurons project to ovBNST

To explore the role of PACAP in regulation of innate fear-related behaviors, we first performed immunostaining for this neuropeptide, focusing on innate fear circuits in mice. We found that PACAP-containing fibers densely innervate the dorsal BNST, but strikingly, specifically only within its oval nucleus (ovBNST) (Fig. 1a, b). Notably, BNST is both structurally and functionally heterogeneous as optogenetic activation of ovBNST was earlier shown to be anxiogenic, whereas activation of the anterodorsal subnucleus (adBNST) was anxiolytic (Kim et al., 2013). Previously, it has been demonstrated that direct infusions of PACAP into BNST result in enhanced innate fear responses (anxiety-like behaviors) (Hammack et al., 2009; Roman et al., 2014), and projections from the basolateral amygdala (BLA) to adBNST were shown to control anxiety (Kim et al., 2013). Thus, the observed pattern of BNST innervation by PACAPergic fibers suggested to us that PACAP may contribute to fear-related behaviors by controlling the flow of signals through BLA-ovBNST-adBNST (BLA-BNST) circuits.

To address this possibility, we first identified the source of PACAPergic projections in ovBNST by injecting a fluorescent retrograde tracer (Katz et al., 1984; Cho et al., 2013) into the mouse ovBNST *in vivo* and performing immunohistochemical labeling for PACAP three weeks later (to allow for the delivery of retrobeads to afferent areas targeting ovBNST) (Fig. 1c). Consistent with a previous report (Missig et al., 2014), many cells in the parabrachial nucleus (PBn) were retrogradely labeled by the fluorescent retrograde beads, indicating that PBn sends direct projections to ovBNST (Fig. 1d). The majority of ovBNST-projecting (fluorescent beads-containing) PBn neurons were immunopositive for PACAP (Fig. 1d-i), suggesting a role for PBn as an endogenous source of this neuropeptide in ovBNST. Neurons in the paraventricular nucleus of the hypothalamus (PVN) were also retrogradely labeled by the fluorescent tracer, but they do not seem to be a source of PACAP in ovBNST, since we did not observe PACAP-immunopositive cell bodies in PVN (not shown). Notably, we found that PACAP’s cognate receptor PAC1 is densely expressed in ovBNST (Fig. 2).

### Photostimulation of PBn fibers results in PACAP release in ovBNST

Using anterograde viral tracing with AAV-hSyn-ChR2-eYFP viral construct, injected into PBn (Fig. 4a), we confirmed the PBn-ovBNST connectivity (Fig. 4b-d). We also confirmed the functionality of ChR2 expression in PBn, observing light-induced photocurrents (Fig. 3b) and spikes (Fig. 3a, c) (under voltage-clamp or current-clamp conditions, respectively) in recorded PBn neurons. To obtain evidence that PACAP could, in fact, be released from PBn fibers, we performed *ex vivo* experiments on slices from mice in which ChR2 was expressed in PBn neurons. We found that photostimulation of ChR2-expressing PBn fibers in ovBNST with a protocol identical to the one that was used in our subsequent behavioral experiments (5-ms pulses were delivered at a 30 Hz frequency for 5 min) resulted in the sustained inward current in ovBNST neurons recorded under voltage-clamp conditions at a holding potential of -70 mV (Fig. 3d) in the presence of antagonists of GABAAR (bicuculline), AMPAR (NBQX), and NMDAR (D-AP5). PACAP exerts its actions through the binding to three G protein-coupled receptor subtypes with similar affinity. One receptor, PAC1, is specific to PACAP, whereas two other receptors, VPAC1 and VPAC2, are shared by PACAP with VIP (Vaudry et al., 2009; Dickson and Finlayson, 2009). Notably, a recent pharmacological study demonstrated that anxiety-promoting effects of PACAP in BNST are mediated by activation of the PAC1 receptor (Roman et al., 2014), and our immunohistochemistry data showed a high level of PAC1 receptor expression in ovBNST (Fig. 2a, b). Thus, we tested the effect of administration of PACAP6-38, known to block PAC1 receptors (Cho et al., 2012; Roman et al., 2014), on the inward current in ovBNST neurons induced by photostimulation of PBn fibers, and found that it was suppressed under conditions of the PAC1 receptor blockade (Fig. 3d-f). To estimate how much PACAP could be released in the ovBNST in the course of photostimulation of PBn projections, we determined a concentration of exogenously-applied PACAP38, a biologically active form of PACAP (Vaudry et al., 2009), producing functional effects identical to those of endogenously-released PACAP. Specifically, we compared peak magnitudes of inward currents which were induced in ovBNST neurons by exogenously-applied PACAP38 to the effects of optogenetic activation of PBn-ovBNST projections. We found that 10 nM PACAP38 induced membrane currents (also blocked by PACAP6-38) in ovBNST neurons identical to those triggered by endogenously-released PACAP (Fig. 3e, f) (PACAP- and photostimulation-induced current amplitudes were 123.5 ± 43.5 pA, *n* = 9 neurons from 4 mice and 118.6 ± 36.7 pA, *n* = 8 neurons from 4 mice, respectively, *P* = 0.93 between groups, Student’s two-tailed unpaired *t-*test). Therefore, PACAP38 in a concentration of 10 nM was used in subsequent *ex vivo* experiments exploring how the signal flow in innate fear circuits is controlled.

**Figure 3.**
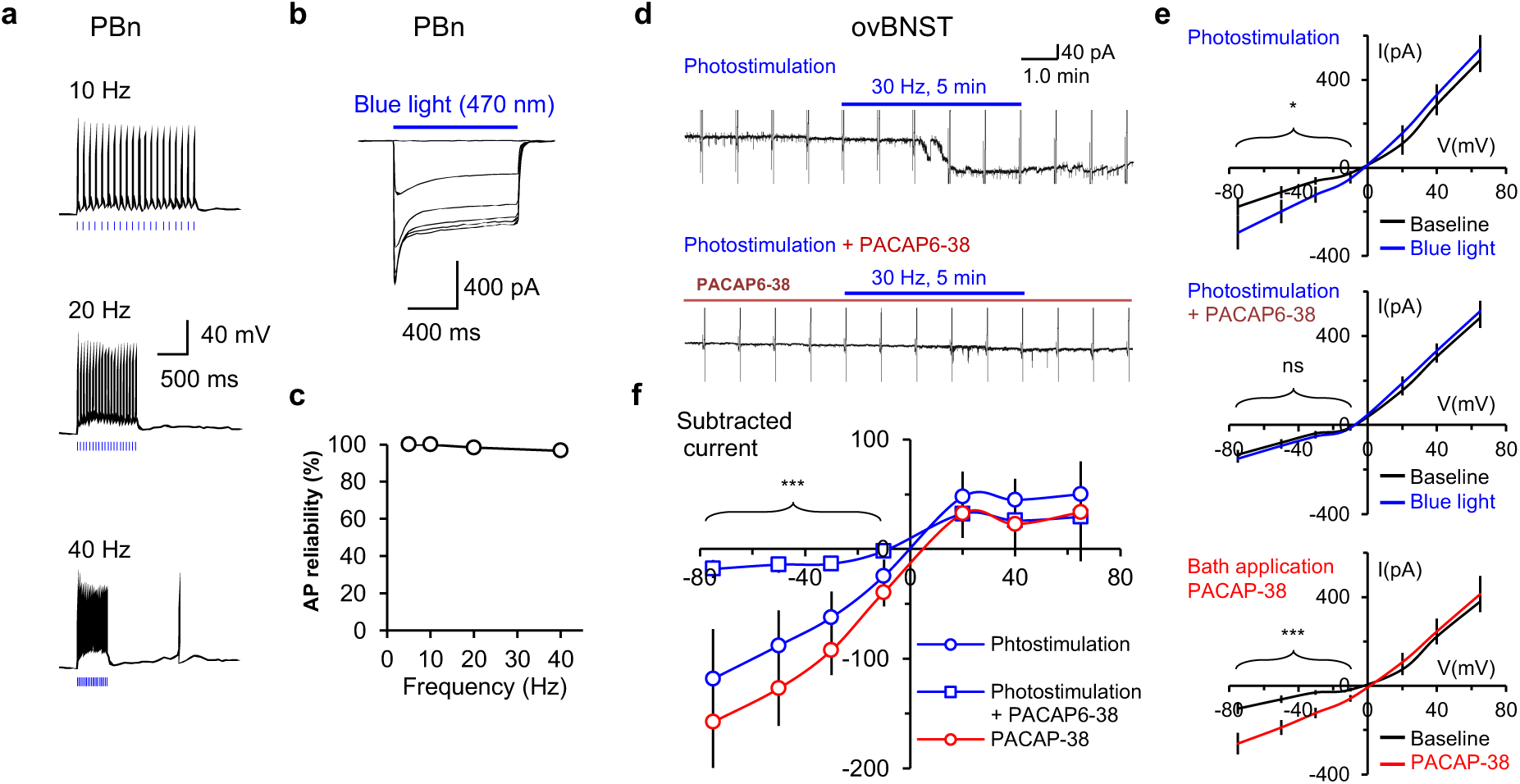
Optogenetic activation of PBn fibers triggers PACAP release in ovBNST. **a**, Action potentials (APs) were triggered in ChR2-expressing PBn neurons in current-clamp mode by 5-ms pulses of blue light at different frequencies (the delivery of pulses is marked by vertical blue bars). **b**, ChR2-mediated photocurrents currents in PBn neurons were evoked by 1-s pulses of 470 nm light (blue horizontal bar) of increasing intensity (0.2 - 1.7 mW/mm^2^). **c**, Plot of AP firing probability versus photostimulation frequency at a 1.7 mW/mm^2^ light intensity (n = 6 neurons). **d**, Top, photostimulation of ChR2-expressing PBn fibers (at blue horizontal bar; 30 Hz, 5 min) resulted in inward current in ovBNST neurons at a holding potential of -70 mV (118.6 ± 36.7 pA, n = 8 neurons from 4 mice). Below, the inward current was suppressed by PAC1 receptor antagonist PACAP6-38 (200 nM). Voltage ramps (3.5-s long, from -75 mV to +65 mV) were applied once a minute during these recordings. The recordings were performed in the presence of NBQX (10 μM), D-AP5 (50 μM), bicuculline (10 μM), TTX (1.0 μM) and CdCl2 (200 μM) (as in Riccio et al., 2009). **e**, Averages of five traces of voltage ramp-induced currents in ovBNST neurons during light-off period (black) and on the peak of photostimulation-induced inward currents (blue) under control conditions (top), in the presence of PAC1 receptor antagonist PACAP6-38 (middle), and in the presence of PACAP-38 (10 nM, PAC1 receptor endogenous agonist; lower). **f**, The baseline-subtracted voltage ramp-induced currents in ovBNST neurons under different conditions. PACAP6-38 (200 nM) suppressed photostimulation-induced voltage ramp-induced membrane currents neurons (19.2 ± 8.6 pA, n = 6 neurons from 3 mice, *P* = 0.04 versus recordings without PAC1R antagonist, Student’s unpaired two-tailed t-test). The current induced by the application of 10 nM PACAP-38 in ovBNST neurons was not different from the current resulting from photostimulation of ChR2-expressing PBn fibers *P* > 0.05). Data are mean ± s.e.m.

**Figure 4.**
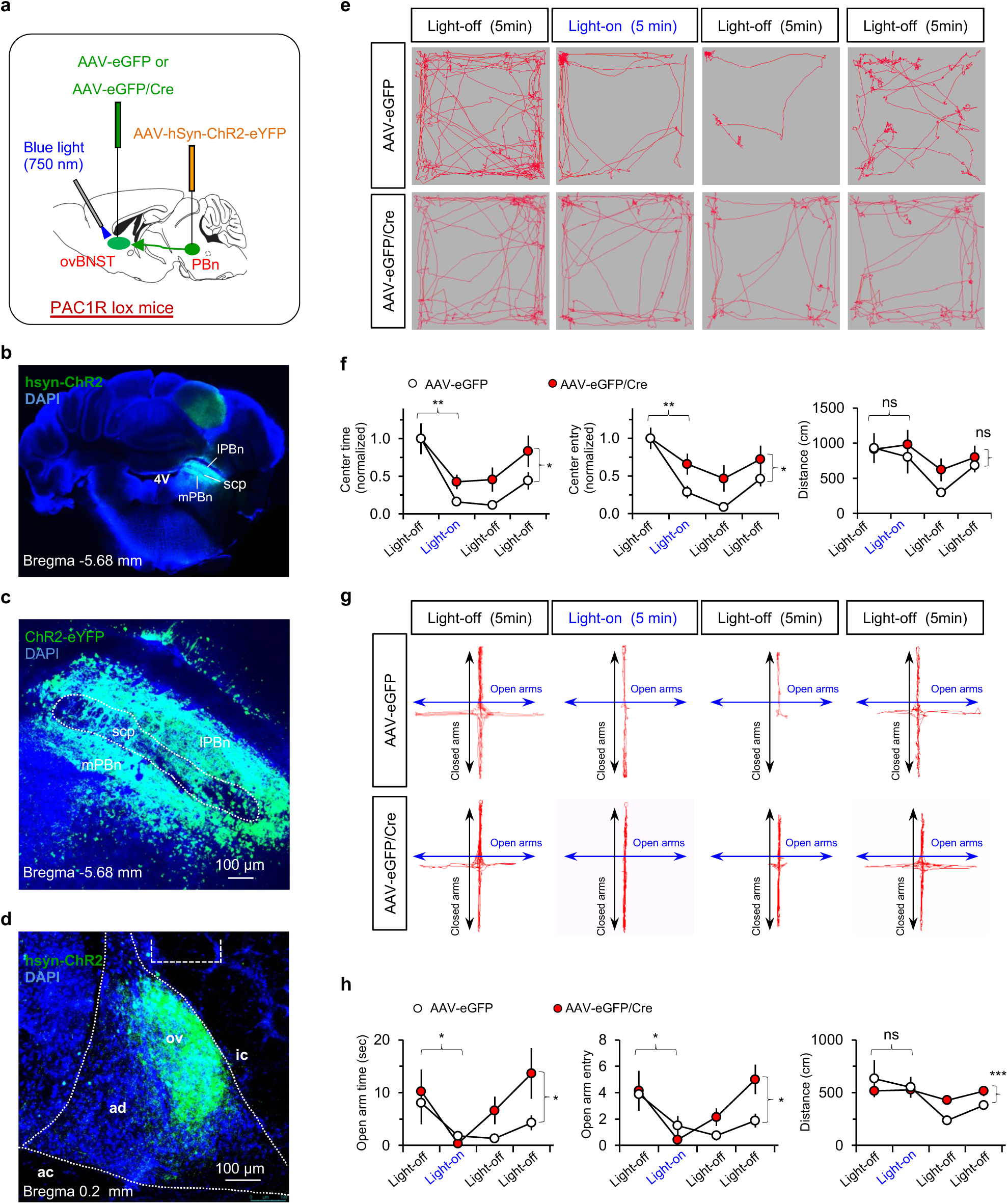
Optogenetic stimulation of PBn-arising fibers in ovBNST enhances anxiety-like behavior through activation of PAC1 receptors. **a**, The experimental design: AAV-hSyn-ChR2-eYFP was injected into PBn in *Adcyap1R1*^fl/fl^ mice to target the PBn-ovBNST projections and PAC1R expression was knocked down in the ovBNST specifically by injecting AAV8.CMV.HI.eGFP-Cre in this subnucleus. **b-d**, Representative confocal images show an injection site of AAV-hSyn-ChR2-eYFP expression in the PBn at low (**b**) and high (**c**) magnification, as well as the ChR2-expressing axons in ovBNST (**d**). lPBn, lateral parabrachial nucleus; mPBn, medial parabrachial nucleus; scp, superior cerebellar peduncle. **e**, Representative behavioral tracks in control mice (top, PAC1R loxP:eGFP) and mice in which PAC1Rs were deleted (bottom, PAC1R loxP:eGFP-Cre) in 20 minutes open field (OF) test consisting of four 5-minutes epochs: light-off epoch (left), light-on epoch (middle-left), light-off epoch (middle-right) and light-off epoch (right). **f**, Summary plots showing the effects of prolonged photostimulation (5 ms-pulses of blue light, at 30 Hz for 5 minutes) of ChR2 expressing fibers in the ovBNST. Both the center time (left) and the center entries (middle) were decreased in control mice (\*\**P* = 0.006 and \*\**P* 0.002, respectively; two-tailed paired *t*-test, n = 7 mice, light off versus light-on) whereas the travel distance was unaffected in control mice (*P* = 0.569), indicating that photostimulation of PBn fibers in ovBNST was anxiogenic. The anxiety-triggering effects of PBn fibers activation in ovBNST were blunted in PAC1R-deleted mice (n = 8 mice) at the level of both the center time (*F*(1,26) = 5.553, *P* = 0.026, two-way ANOVA; AAV-eGFP group versus AAV-eGFP/Cre mice) and center entry (*F*(1,26) = 4.453, *P* = 0.045, two-way ANOVA), whereas there was no difference in the travel distance (*F*(1,26) = 2.866, *P* = 0.102, two-way ANOVA) between the groups. **g**, Representative behavioral tracks in tests designed analogous to **e** but the elevated plus-maze (EPM) test was used instead of OF. **h**, The identical photostimulation of ChR2-expressing PBn fibers in the ovBNST resulted in the reduction of the open arm time (left) and the open arm entries (middle) but not of the travel distance in control mice (*P* = 0.072, 0.071 or 0.338, respectively, paired *t-*test, n = 7 mice, light off versus light-on). In mice with the PAC1R deletion (n =8), anxiogenic effects of PBn fibers stimulation in ovBNST, assayed with the EPM test, were also blunted compared to control (AAV-eGFP-injected mice: open arm time (*F*(1,26) = 5.236, *p* = 0.03, two-way ANOVA) and open arm entries (*F*(1,26) = 7.168, *P* = 0.013, two-way ANOVA). Data are mean ± s.e.m.

### PAC1 receptors in ovBNST mediate anxiety-enhancing effects of PBn-ovBNST projections activation

To test the role of PBn projections to BNST in control of innate fear, we optogenetically stimulated nerve terminals of ChR2-expressing PBn projections in ovBNST of *Adcyap1R1*^fl/fl^ mice in the course of the open field or elevated plus-maze behavioral assays, commonly used to study innate fear-related behaviors (Riccio et al., 2009; Kim et al., 2013). Photostimulation of PBn-arising projections in ovBNST (470 nm, 5-ms pulses at 30 Hz for 5 min) was strongly anxiogenic under control conditions, as mice spent less time in the open field center and entered it less frequently compared to the initial light-off period (Fig. 4a, e, f). Conversely, in the elevated plus-maze test, mice spent less time in the open arms and entered them less frequently upon photostimulation of PBn projections in ovBNST compared to the pre-stimulation baseline (Fig. 4a, g, h). We then performed identically designed behavioral experiments on *Adcyap1R1*^fl/fl^ which received injections of AAV8.CMV.HI.eGFP-Cre into the ovBNST to ablate PAC1 receptors in this BNST subnucleus specifically. The ablation of PAC1R expression in ovBNST resulted in a suppression of innate fear increases triggered by activation of PBn-ovBNST projection (Fig. 4a, e-h). Together, these findings indicate that activation of PBn inputs in ovBNST is strongly anxiogenic and anxiety-driving effects of PBn-ovBNST projections stimulation may be mediated by PAC1 receptors.

### BLA projections to different components of the dorsal BNST are differentially modulated by PACAP

There is evidence that projections from the basolateral amygdala to the dorsal BNST can contribute to control of anxiety states (Kim et al., 2013). To explore whether PACAP modulates the signal flow in BLA-BNST circuits, we first identified targets of BLA projections within the BNST, expressing ChR2-eYFP under the control of the CaMKIIα promoter in BLA neurons. Six weeks later, expressed ChR2 was functionally active and could drive neuronal firing in the BLA in response to photostimulation (as in Cho et al., 2013) (Fig. 5a, b, d-f). ChR2-expressing BLA fibers were found in both the adBNST and ovBNST, though the density of fibers was higher in adBNST (Fig. 5c). Notably, neurons in these two BNST subdivisions did not differ in their inherent membrane properties (Fig. 5g). BLA fibers form functional synaptic contacts on neurons in the BNST, as pulses of blue light (5-ms duration) evoked excitatory postsynaptic currents (EPSCs) in BNST neurons under voltage-clamp conditions at -70 mV. These EPSCs were glutamatergic, as they were blocked by the AMPA/kainate receptor antagonist, NBQX (10 μM) (Fig. 5h, i) and monosynaptic in nature (Fig. 6). Consistent with the observed dense innervation of adBNST by BLA fibers at the light microscopic level, a much greater fraction of adBNST neurons exhibited synaptic responses upon presynaptic photostimulation, and their amplitude was larger in BLA projections to adBNST, compared to inputs to ovBNST (Fig. 5j, k, m, n). Consistent with the finding that PAC1R expression is more abundant in ovBNST compared to adBNST (Fig. 2), exogenously-applied PACAP induced marked potentiation of glutamatergic EPSCs at inputs to ovBNST but had no effect at inputs to adBNST (Fig. 5j, k m, n), indicating that the effects of PACAP on the signal flow in BLA-BNST circuits may be projection specific. The paired-pulse ratio (PPR) of EPSCs, an index of presynaptic function (Zucker and Regehr, 2002), induced by paired presynaptic stimuli at a 50-ms inter-pulse interval was unaffected by PACAP in either of two pathways, indicating that PACAP-induced potentiation at BLA-ovBNST synapses was postsynaptically expressed (Cho et al., 2012) (Fig. 5i, o).

**Figure 5.**
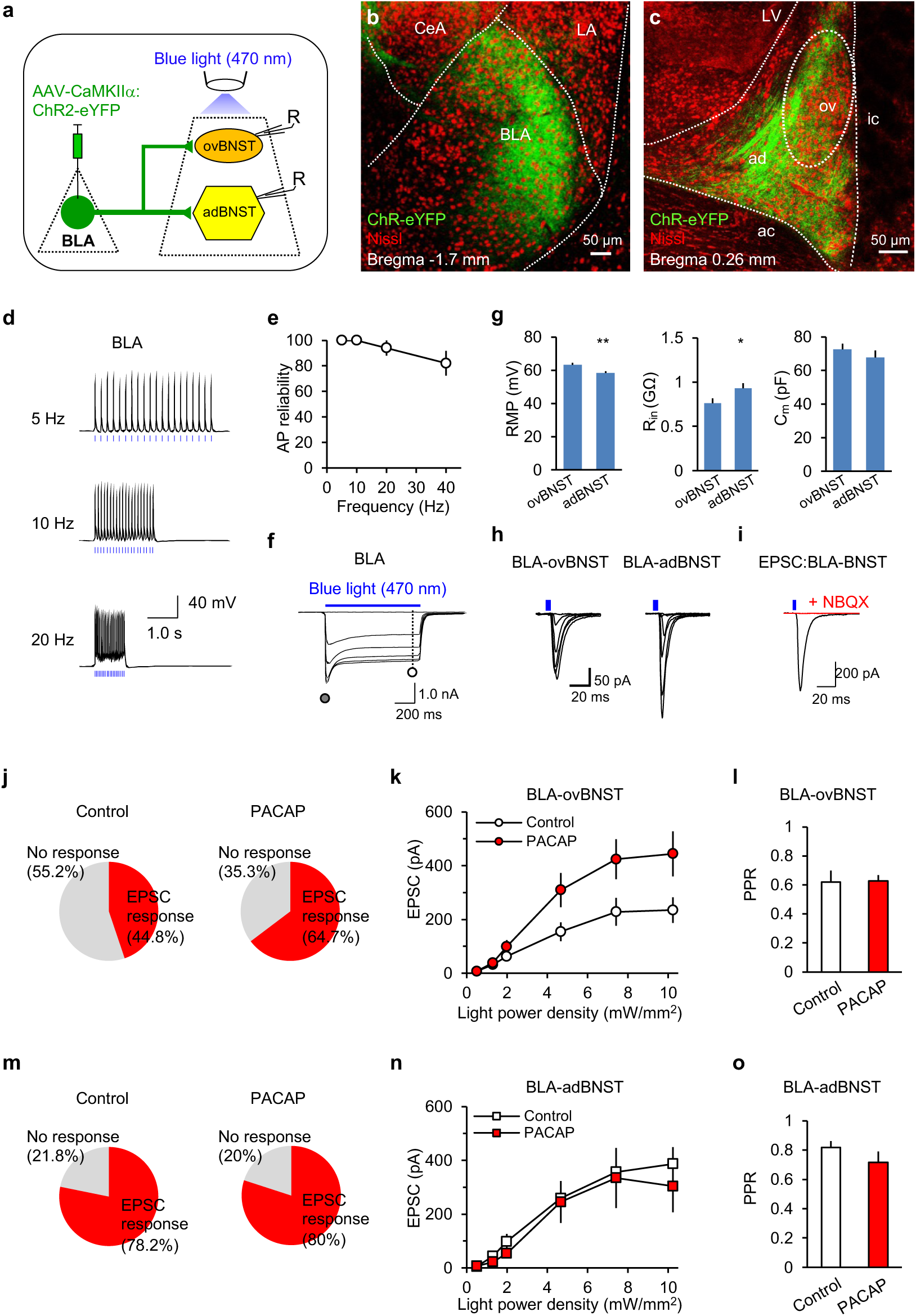
Differential neuropeptide-mediated modulation of synaptic efficacy at distinct BLA projections to the dorsal BNST. **a**, Experimental design for optogenetic projection-specific activation of BLA inputs to ovBNST or adBNST. **b**, Expression of eYFP-ChR2 at the injection site in BLA (green), superimposed with fluorescent Nissl stain (red). LA, lateral nucleus of amygdala; BLA, basolateral nucleus of amygdala; CeA, central nucleus of amygdala. **c**, An image showing ChR2-expressing BLA fibers in the BNST (green). **d**, APs were reliably triggered in BLA neurons in current-clamp mode by 5-ms pulses of blue light at different frequencies (marked by vertical blue bars). **e**, Plot of AP firing probability versus photostimulation frequency at a 1.67 mW/mm^2^ intensity (n = 5 neurons). **f**, ChR2-mediated currents in BLA neurons were evoked by prolonged pulses of 470 nm light (1-s long, blue horizontal bar) of increasing intensity at -80 mV in voltage-clamp recording mode in the presence of NBQX (10 µM), D-AP5 (50 µM) and bicuculline (30 µM). **g**, Resting membrane potential (MP, left), input resistance (Rin, middle) and capacitance (right) in ovBNST (n = 78 neurons) or adBNST (n = 71) neurons. **h**, Light-induced EPSCs recorded from ovBNST (left) or adBNST (right) neurons at a holding potential of -70 mV. **i**, The EPSCs were blocked by glutamate receptor antagonists (10 µM NBQX, and 50 µM D-AP5). **j**, Proportions of neurons in ovBNST which displayed EPSCs in response to photostimulation of BLA fibers under control conditions (left; number of neurons examined was 29) and in the presence of 10 nM PACAP (right; 17 neurons examined). **k**, Input-output curves for peak amplitudes of light-induced EPSCs in the BLA-ovBNST pathway induced by photostimuli of increasing intensity (0.5-10.2 mW/mm^2^) under control conditions (empty symbols; n = 13 neurons from 8 mice) and in the presence of PACAP (10 nM; red symbols; n = 11 neurons from 6 mice; *P* < 0.001 for control versus PACAP, two-way ANOVA). **l**, PACAP had no effect on paired-pulse ratio (PPR) of EPSCs recorded from ovBNST neurons (*P* = 0.92, Student’s unpaired two-tailed *t-*test). Stimulation intensity was 10.2 mW/mm^2^. **m**, Proportions of neurons in adBNST exhibiting EPSCs in response to photostimulation of BLA fibers under control conditions (left; number of neurons examined was 50) and in the presence of 10 nM PACAP (right; 10 neurons examined). **n**, Input-output curves for peak amplitudes of EPSCs in the BLA-adBNST pathway under control conditions (empty symbols; n = 41 neurons from 14 mice) and in the presence of PACAP (10 nM; red symbols; n = 8 neurons from 3 mice; *P* = 0.45 between groups, two-way ANOVA). **o**, PPR of EPSC recorded from adBNST neurons (*P* = 0.26, Student’s unpaired two-tailed *t*-test). Data are mean ± s.e.m.

**Figure 6.**
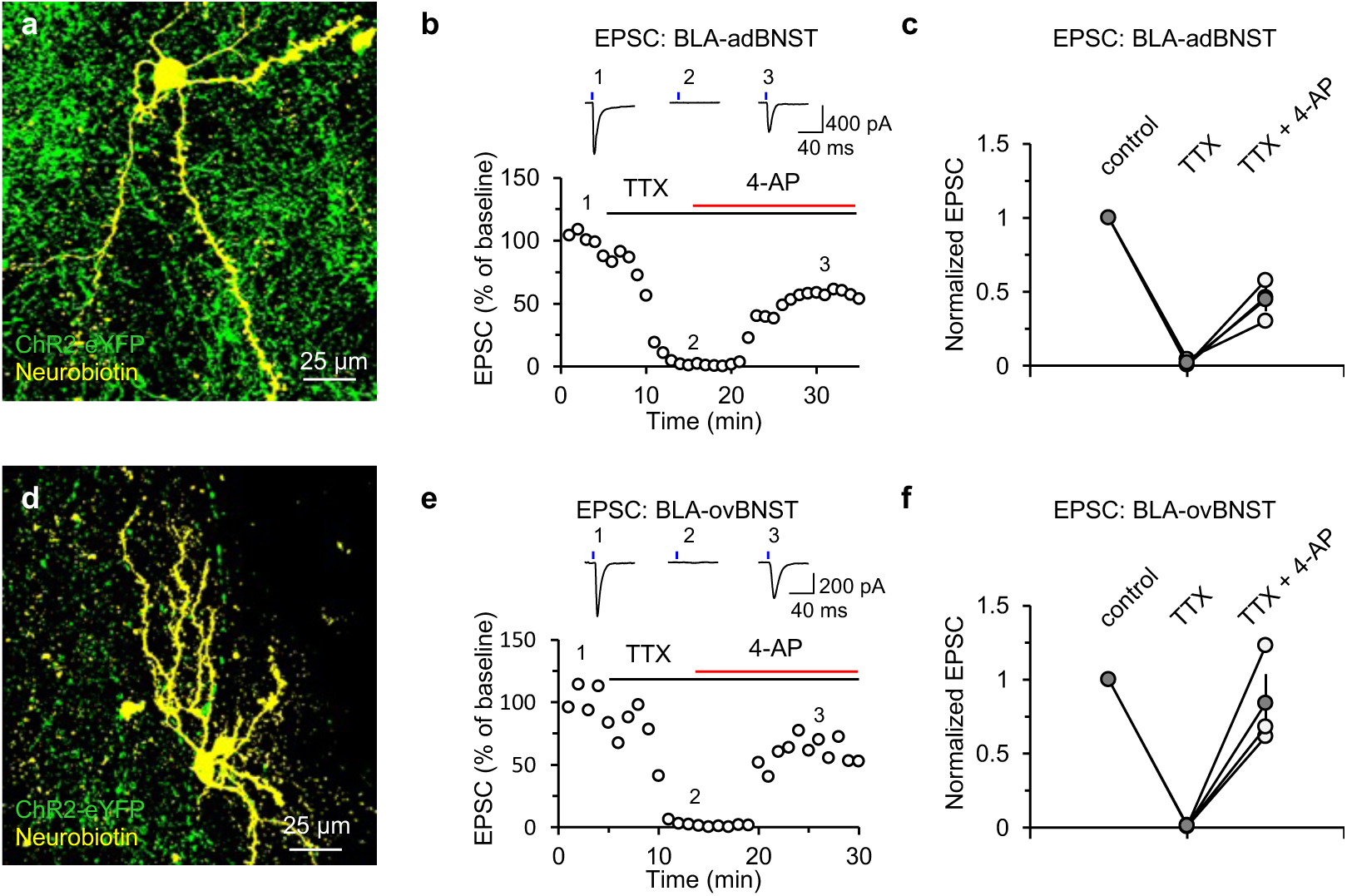
Optogenetically activated BLA inputs to dorsal BNST subdivisions are monosynaptic. **a**, A microscopic image showing intracellularly stained adBNST neuron and ChR2-eYFP-expressing BLA fibers (green fluorescence). Neurons were loaded with the pipette solution containing 6 mM neurobiotin, and visualized with streptavidin, Alexa 568 conjugate. **b**, Rescue of optogenetically-triggered and tetrodotoxin (TTX)-blocked BLA-adBNST EPSCs by 4-aminopyridine (4-AP). Top, representative traces of EPSCs recorded under different experimental conditions. Bottom, EPSCs were evoked by photostimulation of ChR2-expressing BLA fibers and recorded in an adBNST neuron at –70 mV in voltage-clamp mode (1). Bath-applied TTX (1 μM) completely blocked EPSCs (2). Subsequent application of 4-AP (1 mM) in the presence of TTX partially rescued EPSCs (3), indicating monosynaptic nature of light-induced synaptic responses in the BLA-adBNST pathway. EPSC amplitudes were normalized to the baseline mean EPSC amplitude value. **c**, Summary plot showing normalized EPSC amplitudes at BLA-adBNST synapses under three experimental conditions (control, TTX only and TTX + 4- AP). Open circles represent individual experiments whereas filled circles show average values (n = 3 neurons from 3 mice). **d, e**, The experiment was identical to **a-c** but for BLA projections to ovBNST. **f**, Summary plot showing normalized EPSC amplitudes at BLA-ovBNST synapses under three experimental conditions (control, TTX only and TTX + 4-AP). As in **c**, open circles represent individual experiments whereas filled circles show average values (n = 3 neurons from 3 mice). Data are mean ± s.e.m.

### PACAP-mediated increases in synaptically-driven firing of ovBNST neurons result in enhanced feed-forward inhibition in BLA-ovBNST-adBNST pathway

The observed projection-specificity of synaptic effects of PACAP in the BNST, with PACAP-induced strengthening of glutamatergic synaptic projections to ovBNST specifically, indicates that PACAP could selectively modulate synaptically-driven activity of neurons in the oval BNST, possibly increasing their spike output in response to incoming activity, and, therefore, controlling anxiogenesis (Kim et al., 2013). To address these possibilities, we first confirmed that projections from ovBNST to adBNST are purely GABAergic, as focal stimulation of ovBST neurons with minimal intensity electrical pulses triggered inhibitory postsynaptic synaptic currents (IPSCs) in adBNST neurons which were always completely blocked by the GABAA receptor antagonist bicuculline (Fig. 7a-c).

**Figure 7.**
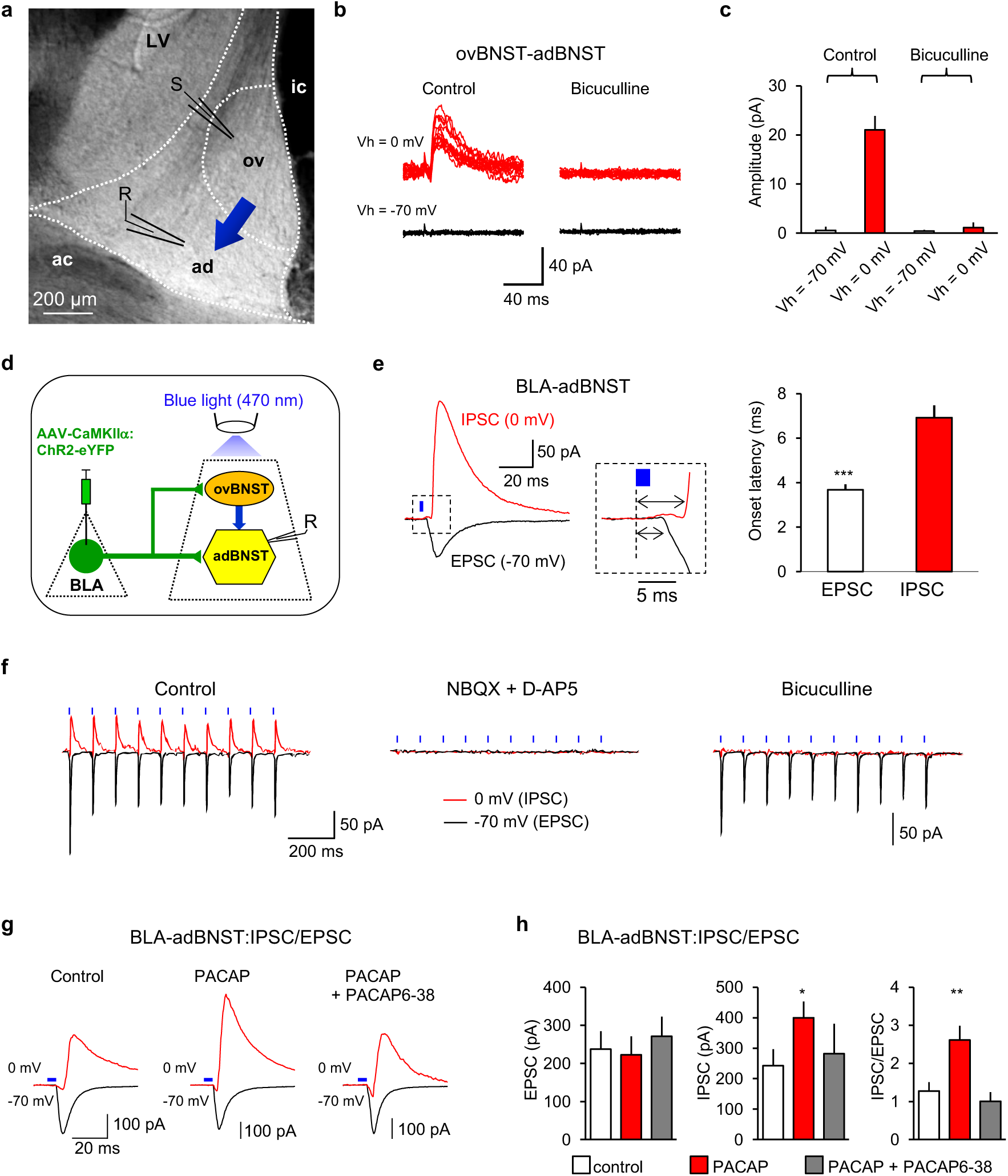
PACAP-mediated enhancement of feed-forward inhibition in BLA-ovBNST-adBNST pathway. **a**, The slice preparation, showing locations of stimulation (S) and recording (R) electrodes in ovBNST and adBNST, respectively. **b**, Synaptic responses were evoked in adBNST neurons by focal stimulation of ovBNST neurons with electrical pulses of minimal intensity at holding potential of -70 mV (black traces, bottom) or 0 mV (red traces, top). Synaptic responses at 0 mV were blocked by the GABAA receptor antagonist bicuculline (30 μM, right), confirming that they were GABAergic IPSCs. **c**, Summary graph showing averaged magnitudes of synaptic responses in control solution (n = 7 neurons) and in the presence of bicuculline (n = 3 neurons), *P* < 0.001 for control versus bicuculline-blocked responses at 0 mV. **d**, A diagram illustrating the neural circuit mediating feed-forward inhibition in BLA-ovBNST-adBNST pathway and experimental design for optogenetic activation of feed-forward IPSCs in adBNST neurons. **e**, Left and middle, synaptic latency analysis for EPSCs and IPSCs recorded in adBNST neurons at -70 mV or 0 mV, respectively. Right: synaptic latency of IPSCs was significantly longer than that of EPSCs (n = 9 adBNST neurons, *P* < 0.001 for EPSC versus IPSC latencies, Student’s paired *t*-test). **f**, Synaptic responses induced by pulses of blue light (5 ms in duration) were recorded in adBNST neurons at holding potentials of –70 mV (black traces) or 0 mV (red traces) under control conditions (left), in the presence of glutamate receptor antagonists (10 µM NBQX and 50 µM D-AP5, middle), and GABAA receptor antagonist (30 µM bicuculline, right). Both glutamatergic and GABAergic responses were blocked by glutamatergic antagonists, indicating the disynaptic nature of GABAAR-mediated IPSCs in adBNST neurons. **g,** Representative traces of synaptic responses in adBNST neurons induced by photostimulation of ChR2-expressing BLA fibers at holding potentials of -70 mV (black trace, bottom) or 0 mV (red trace, top) under control conditions (left), in the presence of PACAP (10 nM, middle) and when PACAP effects were tested with PAC1 receptors blocked by PACAP6-38 (200 nM). **h**, Summary graphs for experiments shown in **i**. PACAP had no effect on the magnitude of BLA-adBNST EPSCs (*P* = 0.81), but significantly enhanced disynaptic IPSCs (*P* = 0.04). It resulted in increased the IPSC/EPSC ratio (*P* = 0.004). The increase was suppressed by PACAP6-38, a PAC1 receptor antagonist (*P* = 0.43 versus control; control group: n = 23 neurons from 12 mice; PACAP: n = 19 neurons from 7 mice; PACAP6-38: n = 11 neurons from 2 mice). Data are mean ± s.e.m. \*\**P*<0.01; \*\*\**P*<0.001.

In ex vivo optogenetic experiments, both glutamatergic EPSCs and GABAergic IPSCs could be recorded in the adBNST neurons in response to photostimulation of ChR2-expressing BLA fibers. Under these conditions, the IPSCs are disynaptic, as their synaptic latency was nearly twice as long compared to EPSCs (Fig. 7d, e) and they are blocked by glutamate receptors antagonists as well as by the GABAA receptor antagonist bicuculine (Fig. 7f), confirming that they are GABAergic. Notably, projections from ovBNST were the source of disynaptic IPSCs in adBNST triggered by photostimulation of BLA-originating projections, since IPSCs in adBNST were suppressed when ovBNST was physically separated from adBNST by cutting connecting fibers. Thus, we recorded EPSCs and IPSCs from the same adBNST neurons, photostimulating BLA fibers, first recording EPSC at a holding potential of -70 mV (close to the reversal potential, Er, for the IPSC), and then recording IPSCs at 0 mV (the Er for AMPAR EPSCs) under control conditions first (Fig. 7g, left), and then in the presence of PACAP (Fig. 7g, center). There was no cross-contamination between EPSCs and IPSCs under these recording conditions (Shin et al., 2006). To determine whether the balance between inhibition and excitation was affected by PACAP, we calculated IPSC/EPSC amplitude ratio under control conditions and in the presence of PACAP. Consistent with our observation that the potentiating effect of PACAP on glutamatergic synaptic transmission was restricted to BLA inputs to ovBNST (Fig. 5k, n), PACAP had no effect on glutamatergic EPSCs at BLA-adBNST synapses in these experiments, but it substantially enhanced the amplitude of disynaptic GABAergic IPSCs in adBNST neurons (Fig. 7g, h). This resulted in increased IPSC/EPSC amplitude ratio, indicating that PACAP enhanced the efficacy of feed-forward inhibition in the BLA-ovBNST-adBNST pathway (Fig. 7h). The potentiating effect of PACAP on feedforward IPSCs in adBNST was blocked by the PAC1 receptor antagonist PACAP6-38 (Fig. 7g, h), providing evidence that this potentiation was mediated by activation of PAC1 receptors. Notably, PACAP had no effect on monosynaptic IPSCs in adBNST neurons induced by direct electrical stimulation of ovBNST (Fig. 8). This is consistent with the notion that PACAP-induced increases in feed-forward inhibition in BLA-ovBNST-adBNST pathway were due to the increased excitatory drive at BLA inputs to ovBNST neurons.

**Figure 8.**
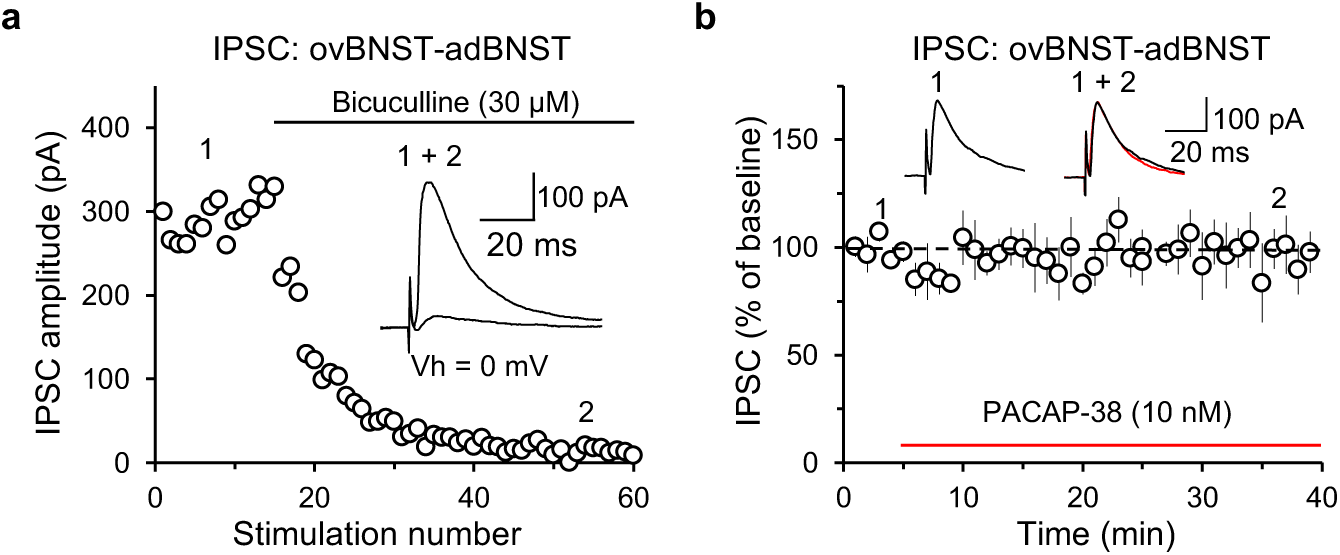
PACAP has no effect on monosynaptic GABAergic IPSCs at ovBNST-adBNST projections. **a**, IPSCs were evoked by a concentric stimulating electrode placed within ovBNST and recorded from adBNST neurons at a holding potential of 0 mV in voltage-clamp mode. The ovBNST-adBNST IPSCs were completely blocked by the GABAA receptor antagonist bicuculline (10 μM). Inset shows averaged traces of IPSCs before and after the application of bicuculline (black horizontal bar). **b**, Addition of PACAP-38 (10 nM) to the external solution did not affect the ovBNST-adBNST IPSC amplitude (n = 6 neurons, *P* = 0.61 for baseline versus PACAP38, Student’s paired *t*-test). The IPSC amplitude was normalized to the baseline IPSC amplitude. Inset shows representative traces of IPSCs recorded before and after PACAP38 application. Data are mean ± s.e.m. \*\*\**P*<0.001.

Under baseline conditions, the differences in the efficacy of excitatory synaptic transmission at inputs to two BNST subdivisions (Fig. 5k, n) were functionally relevant. Thus, the fraction of cells exhibiting synaptically-driven spikes was larger and the probability of spike firing was higher at inputs to adBNST (Fig. 9a-c). To explore whether PACAP-induced changes in synaptic transmission could affect firing output of BNST neurons, we compared the probabilities of extracellular spike firing in neurons in both adBNST and ovBNST recorded in cell-attached mode in response to stimulation of BLA afferents by photo-pulses of increasing intensity under baseline conditions and in the presence of PACAP. PACAP-induced enhancements of excitatory synaptic efficacy in BLA projections to the ovBNST (as shown in Fig. 5j, k, m, n) resulted in the increased spike output of ovBNST neurons driven by activation of BLA fibers (Fig. 9d). The latter is consistent with the PACAP-induced shift in inhibition/excitation balance in the adBNST during activation of BLA-originating projections toward greater functional efficiency of inhibition (as ovBNST, also activated by BLA inputs, sends GABAergic projections to adBNST; Kim et al., 2013; Fig. 7 in this study). Strikingly, synaptically-driven firing output of ovBNST and adBNST neurons was modified by PACAP in opposite directions: it was increased in ovBNST, but decreased in adBNST (Fig. 9d, e). This finding suggests a scenario under which PACAP selectively modulates synaptically-driven activity of neurons in the ovBNST, increasing their spiking in response to incoming activity, and, therefore, inhibiting adBNST and decreasing its synaptically-driven spike output. The decrease in firing output of adBNST neurons may explain why PACAP in BNST is anxiogenic, since optogenetic inhibition of adBNST was shown to enhance anxiety-like behaviors (Kim et al., 2013).

**Figure 9.**
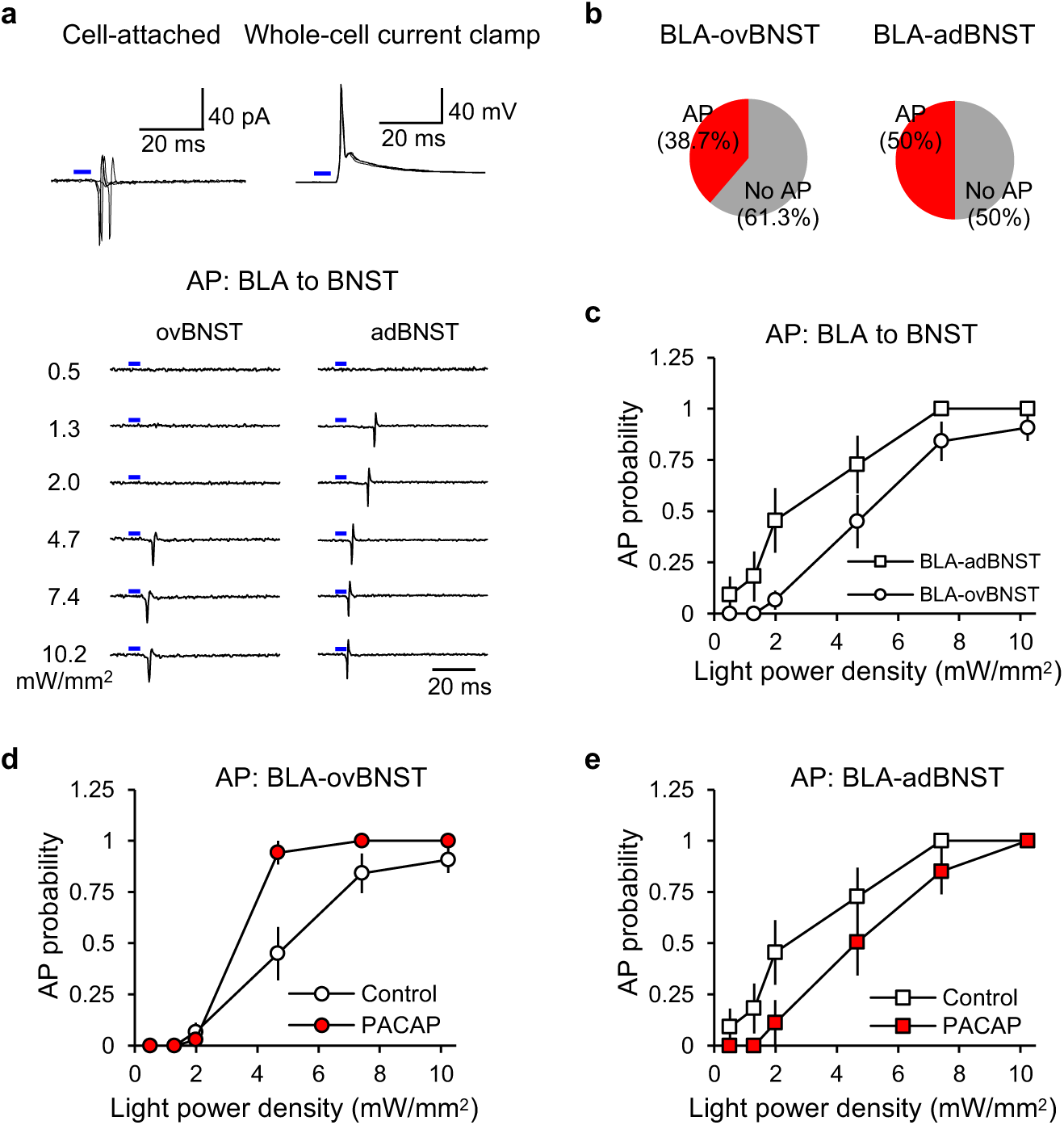
PACAP alters probability of synaptically-driven spike output in BLA-adBNST and BLA-ovBNST projections in opposite directions. **a**, Top, superimposed synaptically-driven extracellular spikes evoked in an adBNST neuron by photostimulation (5-ms pulses) of ChR2-expressing BLA fibers and recorded in cell-attached patch configuration (left). Right, recordings under current-clamp conditions from the same adBNST cell after establishing a whole-cell configuration. Bottom, examples of responses recorded in ovBNST (left) or adBNST (right) neurons in a cell-attached patch configuration in a control external solution to photostimulation pulses of increasing intensity (stimulation light intensities were in a range of 0.5–10.2 mW/mm^2^). **b**, Proportion of neurons in ovBNST (left; n = 22 neurons from 12 mice) or adBNST (right; n = 24 neurons from 9 mice) which displayed extracellular spikes in response to photostimulation of BLA fibers. **c**, Probability of AP firing induced by photostimulation of BLA projections. Under control conditions, the probability of synaptically-driven AP firing as a function of photostimulation intensity was significantly higher at BLA inputs to neurons in adBNST compared to ovBNST (for ovBNST recordings, n = 12 neurons from 6 mice; for adBNST recordings, n = 11 neurons from 7 mice; *F*1,125 = 14.47, *P* < 0.001, two-way ANOVA). **d**, The probability of synaptically-driven AP firing was increased in BLA-ovBNST projections by PACAP (10 nM; *F*1,102 = 8.36, *P* = 0.005 for control conditions versus PACAP, two-way ANOVA; n = 12 neurons from 6 mice and n = 7 neurons from 6 mice for control and PACAP treatment groups, respectively). **e**, PACAP decreased the probability of synaptically-driven AP firing in BLA-adBNST projections (*F*1,108 = 7.77, *P* = 0.006 for control conditions versus PACAP, two-way ANOVA; n = 11 neurons from 7 mice and n = 9 neurons from 5 mice for control and PACAP treatment groups, respectively). Graphs illustrating AP firing in ovBNST (in **d**) or adBNST (in **e**) neurons in control external solution are the same as in **c** for comparison. Data are mean ± s.e.m.

### Repeated stress-triggered anxiety enhancements are associated with plastic changes in the BLA-ovBNST-adBNST circuits identical to PACAP-induced synaptic effects

What might the conditions under which PACAP released endogenously could contribute to behavioral mechanisms? Notably, chronic (repeated) stress is anxiogenic and is associated with increased expression of both PACAP and PAC1 receptors in the BNST (Roman et al., 2009). Thus, we explored repeated stress-induced changes in the signal flow in innate fear circuits, focusing on the mechanisms of neurotransmission in BLA-BNST circuits, affected by PACAP (see above). In these experiments, we expressed ChR2-eYFP under the control of the CaMKIIα promoter in BLA neurons. 6 weeks after the surgery, mice received repeated footshocks (0.75 mA, 3 s-long; twice a day with 1-min inter-trial interval) for 7 days (Fig. 10a). Freezing behavior was assayed during a 5-min-long period prior to the delivery of shocks and subsequently quantified in the “shock context” on each following day to assess the increase in stress-induced fear (Fig. 10b). On day 8, mice were tested in the open field test (OF) to probe the effect of repeated stress on anxiety levels. We found that repeated stress was anxiogenic under our experimental conditions (Fig. 10c). To evaluate the effects of stress on synaptic transmission in BLA-ovBNST projections, we recorded excitatory postsynaptic currents at BLA-ovBNST synapses in ovBNST neurons while ChR2-expressing BLA fibers were stimulated by pulses of blue light (5- ms duration). Consistent with our earlier observations, these EPSCs were glutamatergic and were blocked by the AMPA/kainate receptor antagonist, NBQX (10 μM), and were monosynaptic in nature. The synaptic efficacy in BLA-ovBNST projections, assayed with the input-output curves for light-induced EPSCs obtained in response to photostimuli of increasing intensity, was significantly enhanced in mice which received repeated footshocks compared to control mice (Fig. 10f). The strengthening of synaptic transmission was not associated with changes in paired-pulse ratio (PPR; 50-ms inter-pulse interval) and thus was postsynaptic in origin (Fig. 10g). Consistent with this notion, we found that the AMPAR/NMDAR EPSC amplitude ratio was increased in repeatedly stressed mice. To obtain values of the AMPAR/NMDAR EPSC amplitude ratio, the amplitude the AMPA receptor EPSC recorded at -70 mV under whole-cell voltage-clamp condition was divided by the amplitude of the NMDAR EPSC recorded at +40 mV and measured 40 ms after the peak of the EPSC at -70 mV. Notably, repeated stress was associated with significant increases in both PACAP and PAC1 receptor immunoreactivity in ovBNST, indicating that PACAP and PAC1R expression were increased in repeatedly stressed mice (Fig. 10i-l). We also recorded both photostimulation-induced glutamatergic EPSCs and GABAergic inhibitory postsynaptic currents (IPSCs) in same adBNST neurons (as in Fig. 7) in slices from control and repeatedly stressed mice. We confirmed that the IPSCs were disynaptic as their synaptic latency was nearly twice as long compared to EPSCs and they were blocked by glutamate receptors antagonists. Thus, we recorded EPSCs and IPSCs from the same adBNST neurons, photostimulating BLA fibers, first recording EPSC at a holding potential of -70 mV (close to the reversal potential, Er, for the IPSC), and then recording IPSCs at 0 mV (the Er for AMPAR EPSCs), in slices from control mice and mice which received footshocks repeatedly. Consistent with our findings with exogenously-applied PACAP, we found that the IPSC/EPSC amplitude ratio was significantly increased in repeatedly stressed mice compared to control animals (Fig. 10h). The latter results indicates that repeated stress results in inhibition of adBNST through the mechanisms analogous to those observed with neuropeptide PACAP, suggesting that repeated stress- and PACAP-induced changes in neurotransmission on BLA-ovBNST-adBNST circuits may share common synaptic mechanisms.

**Figure 10.**
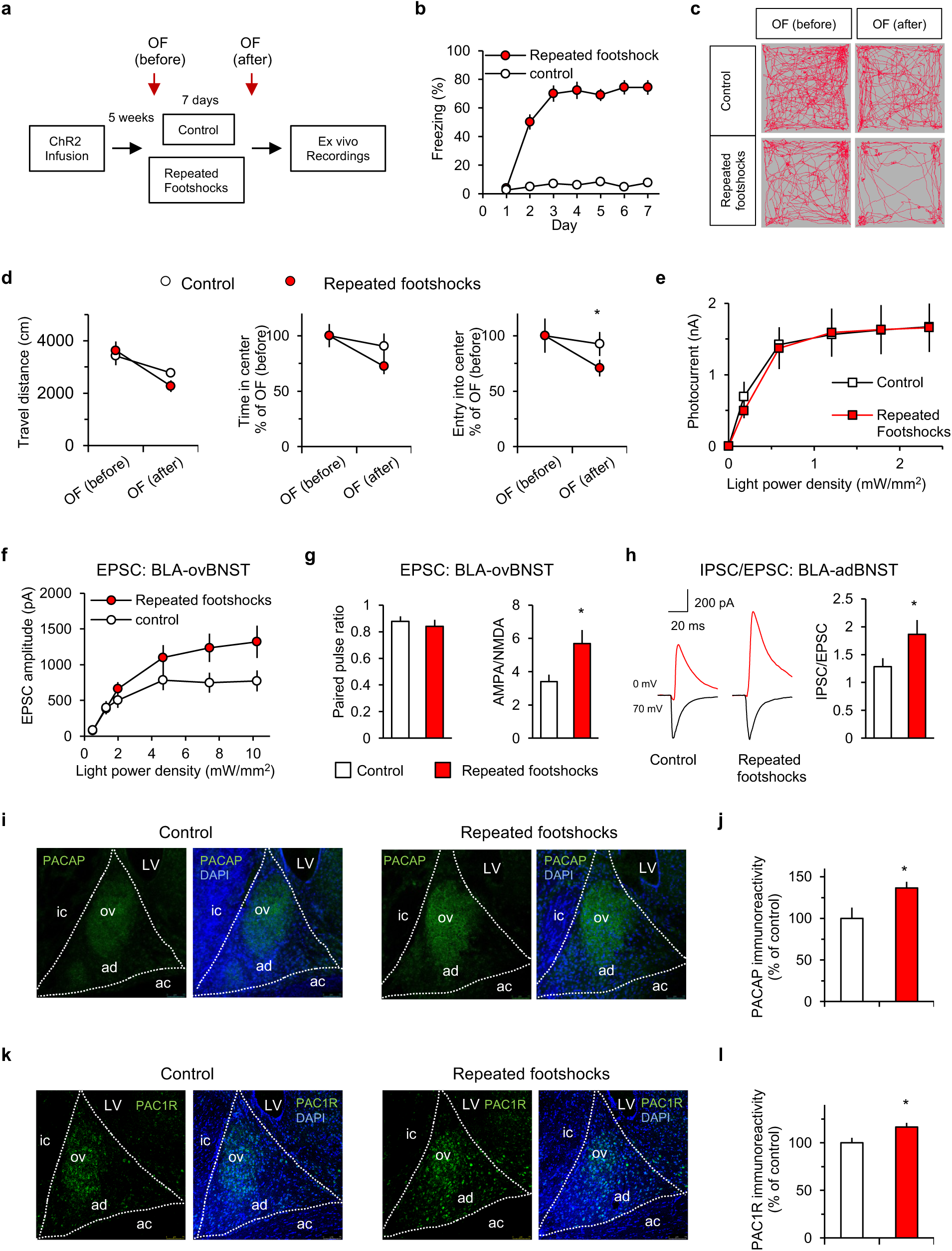
Anxiety-inducing repeated stress is associated with increased feed-forward inhibition at BLA-ovBNST-adBNST projections and enhanced PACAP signaling in ovBNST. **a**, Experimental design for examining the effects of repeated stress (footshocks) on anxiety levels and BLA-BNST circuit function. Mice receiving repeated foot shock were exposed to the 7-day repeated stress paradigm in a plexiglas conditioning chamber (Med Associates, St. Albans, VA) 30 x 25 x 35 cm (L x W x H). After a 5 minutes acclimation period, two 0.75 mA 3 second scrambled footshocks were delivered through the grid floor with a 1 min inter-trial interval (Hammack et al, 2009; Roman et al, 2012). Control mice were placed in the conditioning chamber for the same period of time without foot shocks. **b**, Repeated footshocks results in higher freezing in experimental animals during the 7-day-long behavioral procedure (n = 14 control mice and 14 repeatedly shocked mice, *F*(1,175) = 649.8, *P* < 0.001, two-way ANOVA). **c**, Examples of behavioral tracks in open field tests before (top two panels) and after (bottom two panels) 7-day foot-shock protocols in two groups of animals. Note a marked decrease in time spent in the center in a mouse which received footshocks. **d**, Summary plot of the travel distance (left), time spent in the center (middle), and entry into center (right, *P* = 0.039, unpaired *t*-test) in the second OF test between control mice (n = 11) and repeatedly stressed mice (n = 11). **e**, Summary plot of the amplitude of photocurrents induced by the 1-s pulse of blue light as a function of increasing light intensity in control mice (black open square, n = 10 cells from 4 mice) and repeatedly stressed mice (red open square, n = 10 from 5 mice, *F*(1,108) = 0.078, *P* = 0.78, two-way ANOVA). **f**, Input-output curves for peak amplitudes of light-induced EPSCs in the BLA-ovBNST pathway induced by photostimuli of increasing intensity (0.5-10.2 mW/mm^2^) were different between control mice (empty symbols; n = 25 neurons) and the mice which received footshocks for 7-days (red symbols; n = 22 neurons; *F*(1,285) = 284, *P* = 0.002, two-way ANOVA). **g**, Repeated footshocks resulted in the increased AMPA/NMDA EPSC amplitude ratio (right, n = 13 cells in control mice and 22 cells in repeatedly stressed mice, *P* = 0.92, Student’s two-tail unpaired *t*-test) but had no effect on paired-pulse ratio (PPR) of EPSCs recorded in ovBNST neurons (left, n = 28 cells in both control and footshocked mice, *P* = 0.747, Student’s two-tailed unpaired *t*-test). The stimulation intensity in PPR experiments was 10.2 mW/mm^2^. **h**, Left: Representative traces of synaptic responses in adBNST neurons induced by photostimulation of ChR2-expressing BLA fibers at holding potentials of -70 mV (black trace, bottom) or 0 mV (red trace, top) in control mice (left) and mice receiving repeated footshocks (right). Right: summary plot of IPSC/EPSC ratio at BLA-adBNST projections was increased in mice receiving repeated footshocks (n = 29 neurons in control mice and 20 neurons in mice which received footshocks for 7 days, *P* = 0.044, two-tailed unpaired *t*-test). **i**, Representative confocal images showing immunohistological staining for PACAP (DAPI counterstained) in BNST in control mice (left) and repeatedly stressed mice (right). **j**, PACAP immunoreactivity, estimated by measuring PACAPergic fluorescence intensity in ovBNST, was increased in repeatedly stressed mice (*P* = 0.011, 5 control mice versus 5 footshocked mice, two-tailed unpaired *t*-test). **k**, Representative confocal images showing immunohistological stainings for PAC1R. **l**, PAC1R immunoreactivity in the ovBNST was increased in repeatedly stressed mice (*P* = 0.013, 5 control mice versus 5 footshocked mice, two-tailed unpaired *t*-test).

## Discussion

Our results provide evidence that neuropeptide PACAP, released in the ovBNST from PBn-arising projecting fibers, exerts its actions on the signal flow in anxiety-driving microcircuits through the induction of LTP-like potentiation of glutamatergic synaptic transmission at BLA-ovBNST synapses, not through the direct actions on the functional properties of ovBNST neurons (e.g., changes in the membrane polarization or input resistance). This is a key finding explaining how the neuropeptide-mediated modifications in the function of BLA-ovBNST-adBNST circuits may trigger anxiogenesis. Whereas it has been demonstrated earlier that direct infusions of PACAP into BNST trigger anxiety-like behavioral responses (Hammack et al., 2009; Roman et al., 2014), synaptic and network mechanisms of PACAP-mediated modulation of the signal flow in neural circuits, contributing to behavioral manifestations of anxiety, have not been previously studied and remained unknown. To our knowledge, there was only one study in which the contribution of BNST to control of anxiety was explored at the neural circuit level (Kim et al., 2013). However, the role of BLA-ovBNST-adBNST pathway in these mechanisms was not explored in the mentioned work. This is an important issue, which we now addressed, as our findings show that this pathway plays an essential role in control of anxiety states. In our experiments, the behavioral effects of PBn afferents’ stimulation in the ovBNST were due to the release of PACAP, as the ablation of PAC1Rs in ovBNST in PAC1 receptor “floxed” mice by injecting the *AAV-GFP/Cre* construct into this BNST subdivision blunted the anxiety-producing effects observed when PBn-ovBNST projections were optogenetically activated. This provides evidence that PACAP/PAC1R signaling in ovBNST may contribute to control of anxiety in isolation from other neuromediator systems in the course of anxiety-probing behavioral tests.

What might be the molecular mechanisms possibly mediating PACAP-induced potentiation of synaptic transmission at BLA-ovBNST synapses? We have previously published a study in which we showed that PACAP may induce LTP-like synaptic strengthening at BLA-CeL projections through a postsynaptic mechanism involving enhanced synaptic targeting of GluR1 subunit-containing AMPA receptors (Cho et al., 2012). The analogous mechanisms could underlie PACAP-triggered potentiation at glutamatergic BLA inputs to ovBNST neurons. Interestingly, the anxiogenic effect of PBn-arising fibers stimulation in ovBNST was maintained after the first light-on period in anxiety-probing behavioral tests. This is consistent with the notion that endogenously-released PACAP may induces LTP-like synaptic enhancements in BLA-ovBNST projections.

Notably, in addition to PBn, we observed ovBNST-deposited retrobeads in the BLA. This is not surprising given that BLA sends strong projections to ovBNST. Moreover, we found retrobeads in the medial division of the central nucleus (CeM). However, neither BLA or CeM were found to contain PACAP-positive neurons. We demonstrated previously that CeL (lateral division of the CeA) is densely innervated by PACAPergic fibers (Cho et al., 2012). As CeL sends GABAergic projections to CeM, which, in turn, projects to ovBNST, PACAP-induced potentiation of excitatory inputs from BLA to CeL may result in enhanced inhibition of CeM and, thus, disinhibition of ovBNST (because CeM neurons are also GABAeric), thus potentially contributing to the mechanisms of anxiogenesis. It might be interesting to address these possibilities in future studies. Retrobeads were also observed in the paraventricular nucleus of the hypothalamus (PVN). However, we did not observe PACAP-positive cells in the PVN, indicating that PVN is not the source of PACAP in ovBNST.

We demonstrated also that PACAP-dependent neuromodulation of the studied circuits is implicated in the mechanisms of behavioral defense during specific environmental conditions, such as repeated (chronic) stress. Specifically, our data suggest that under non-anxiogenic conditions, the level of innate fear (i.e., anxiety) might be kept low through maintaining adBNST activation by projections from BLA, which under baseline conditions are stronger than BLA projections to ovBNST (Fig. 11). However, the release of PACAP from PBn fibers in BNST under anxiety-provoking conditions (Hammack et al., 2015), resulting in potentiation of neurotransmission at BLA-ovBNST synapses, would lead to the functional recruitment of ovBNST, inhibiting adBNST and its spike output to the target areas (e.g., lateral hypothalamus), and result in increased anxiety-related behaviors (Fig. 11). Accordingly, our work reveals how anxiety could be controlled by neuropeptides (PACAP, specifically) at the level of underlying neural circuits, by inducing changes in the signal flow between their different structural components. The reported findings may potentially provide clues to understanding the mechanisms of anxiety disorders, reflecting dysregulation of communication between different components of the neural circuits underlying innate fear.

**Figure 11.**
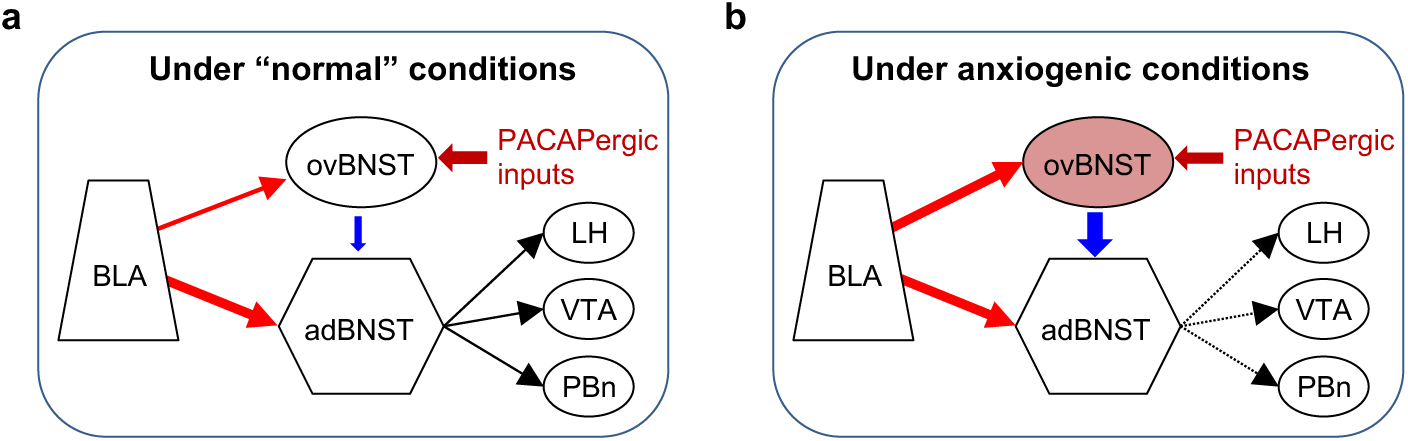
Modulatory roles of PACAP in the function of anxiety-controlling neural circuits. **a,** A diagram illustrating the signal flow in BLA-BNST circuits under “normal” (non-anxiogenic) conditions when excitatory BLA inputs to adBNST are much stronger than projections to ovBNST. This would result in maintaining a “low” anxiety level as excitation of adBNST suppresses behavioral responses to innately fearful stimuli (Kim et al., 2013). **b**, Under anxiety-provoking conditions (e.g., chronic stress), endogenously released PACAP (which is mostly released in ovBNST) would induce LTP-like enhancements in BLA-ovBNST projections (but not at BLA inputs to adBNST), resulting in a functional recruitment of ovBNST. The latter would lead to inhibition of adBNST neurons, as ovBNST cells are GABAergic and they project to adBNST. Through these mechanisms, the spiking output of adBNST neurons will be diminished, thus driving anxiogenesis (Kim et al., 2013). Red and blue arrows indicate glutamatergic and GABAergic projections, respectively, and changes in the thickness of lines indicate changes in synaptic efficacy. PACAPergic input is provided by projections from the PBn. Black arrows show projections to the downstream structures implicated in physiological manifestations of innate fear responses LH, lateral hypothalamus; VTA, ventral tegmental area.

## Abbreviations

PACAP: pituitary adenylate cyclase-activating polypeptide
BLA: basolateral amygdala
BNST: bed nucleus of the stria terminalis
ovBNST: oval BNST
adBNST: anterodorsal BNST
PBn: parabrachial nucleus (PBn)
Scp: superior cerebellar peduncle
CeA: central nucleus of amygdala
EPSCs: excitatory postsynaptic currents
IPSCs: inhibitory postsynaptic currents
4-AP: 4-aminopyridine
AMPAR: α-amino-3-hydroxy-5-methyl-4-isoxazolepropionic acid receptor
NMDAR: N-methyl-D-aspartate receptor
GABA: γ-Aminobutyric acid
LTP: long-term potentiation

## Acknowledgements

We thank Kyung-Jun Park for help and constructive discussions. This work was supported by grants P50MH115874 (to KJR and WAC); R21MH108022 (to VYB) R01MH123993 (to VYB); R01MH108665 (to KJR and VYB). RA was supported by a NARSAD Young Investigator Grant (#22434) and the Ramón y Cajal Program (RYC-2014-15784).

